# Genome-Scale reconstruction of *Paenarthrobacter aurescens* TC1 metabolic model towards the study of atrazine bioremediation

**DOI:** 10.1101/536011

**Authors:** Shany Ofaim, Raphy Zarecki, Seema Porob, Daniella Gat, Tamar Lahav, Yechezkel Kashi, Radi Aly, Hanan Eizenberg, Zeev Ronen, Shiri Freilich

## Abstract

Atrazine is an herbicide and pollutant of great environmental concern that is naturally biodegraded by microbial communities. *Paenarthrobacter aurescens* TC1 is one of the most studied degraders of this herbicide. Here, we developed a genome scale metabolic model for *P. aurescens* TC1, *i*RZ1176, in order to study the atrazine degradation process and mechanism at organism level. Constraint based flux balance analysis and time dependent simulations were used to explore the bacteria’s phenotypic landscape. Simulations led to model based predictions for growth and atrazine degradation efficiency in myriad of media supplemented with different combinations of carbon and nitrogen sources. Simulations aimed at designing media optimized for supporting growth and enhancing degradation, bypassing the need in strain design *via* genetic modifications. Predictions were tested *in vitro* and experimental growth and degradation measurements support and validate the reliability of the model. Model simulations have prioritized degradation and growth enhancement effect of various amino acid and co-factors. The model presented here can potentially serve as a predictive tool towards achieving optimal biodegradation efficiencies and for the development of ecologically friendly solutions for pollutant degradation.

## Introduction

Atrazine (2-chloro-4-ethylamino-6-isopropylamino-1,3,5-triazine) is an herbicide and pollutant of great environmental concern. Its residues are found in soil samples decades after it was last applied and were shown to chronically leach into local aquifers^1^. Areas contaminated with hazardous materials, including atrazine, are rapidly increasing worldwide introducing a need in remediation approaches. Bioremediation - an environmental bioprocess in which naturally occurring organisms are used for breaking down hazardous substances into less toxic or non-toxic substances - is increasingly acknowledged as a cost-effective strategy for environmental cleaning^2–4^. Critical environmental bioprocesses are naturally carried out by bacteria and are related to removal of pollutants from water, soil or air^5^. Thus, bacterial bioremediation is widely applied for the degradation of various organic pollutants ^6–9^ including herbicides^9–13^. Bioremediation may occur naturally or through the stimulating addition of fertilizers, electron acceptors, etc. These amendments encourage the growth of the degrading microbes within the medium or environment in a process termed biostimulation ^3,14^.

One of the well-known atrazine degraders is the gram positive *Paenarthrobacter aurescens* TC1^15–19^. *P. aurescense* TC1 is described as nutritionally versatile bacterium that is capable of utilizing a wide array of complex metabolites, including agriculturally used herbicides^4,20,21^. The full hydrolytic pathway for atrazine degradation was reported and described in *P. aurescens* TC1^21^. Moreover, This bacterium was previously reported to biodegrade atrazine much more efficiently than most atrazine degrading bacteria including *Pseudomonas* sp. ADP ^22^. Intermediates, created in the degradation process are further metabolized by the organism and can provide sole sources for carbon and nitrogen^18^. While *P.*ADP can only use the hydrolytic products of the ring cleavage as nitrogen but not as carbon; TC1 can use the side groups of atrazine as source for carbon, nitrogen and energy consuming up to 3,000 mg of atrazine per liter ^18^. As such, *P. aurescens* TC1 is an ideal candidate for the study of atrazine bioremediation using a genome scale metabolic model reconstruction.

Modern tools of genomics, transcriptomics, metabolomics, proteomics, signaling systems and synthetic biology have opened new avenues for biotechnological advances and are increasingly applied for promoting better understanding of complex biological systems. In recent years, constraint based metabolic modelling approaches have become widely used as an *in silico* tool for organism-level phenotyping and the subsequent development of metabolic engineering strategies ^23,24^. Generally, such approaches follow four key steps: (1) data acquisition – mainly genome sequencing information, basic cell-physiological and biochemical knowledge and some experimental data on cell growth; (2) model reconstruction - translating data into structured mathematical representation; (3) constraint-based optimization simulations - the prediction of growth rate, substrate uptake rates, and byproduct rates under different growth conditions or following knockout mutations, in the absence of kinetic information^25–31^; and (4) experimental validation. The potential of such models for the investigation of optimal processing is now acknowledged and practiced^5,32–35^. Examples for applicative use include the optimal production or utilization of industrial compounds such as xylose^36^, biofuels ^37^,vitamins ^38^ and drug development^39^. In bioremediation, genome scale metabolic modeling approaches were applied for the design of *Geobacter sulfurreducens* strain capable of increased electron transfer and a higher Fe(III) respiratory that was beneficial for environmental bioremediation^40^. Application of genome scale metabolic modeling for strain design is often based on the Optknock algorithm ^41^ and its derivatives which search for the optimal gene knockouts for a desired metabolic production. However, unlike industrial use of engineered strain for enhanced production in monoculture, introduction of exogenous species into natural habitat is far from trivial and hampers the application of modeling approaches towards bioremediation solutions. An alternative strategy is the use of modeling for the design of optimal conditions that will enhance processes (degradation or synthesis) carried by the endogenous community^42^. For example, interactions between *Geobacter* and *Rhodoferax* – a competitor Fe(III)-reducers in anoxic subsurface environments - were captured and described using metabolic modelling and allowed identification of optimal degradation conditions^43^. Models can also predict the benefits of cooperative and commensal interactions to degradation efficiency. Xu *et al* have demonstrate the use of metabolic models for predicting interactions between *P.aurescens* TC1 and endogenous soil species. Exchange metabolites secreted by non-degraders species contribute for enhancing *P.aurescens* TC1 growth and expediting degradation efficiency^44,45^. Here, we directly focused on modeling-based identification of media supplements that can enhance degradation. We describe the process of manual curation of *P.aurescens* TC1 draft metabolic model, its application for predicting growth and atrazine degradation in media containing different carbon and nitrogen sources based on a dynamic (capable of reporting changes over time) and non-dynamic flux based algorithms, and the experimental testing of the predictions.

## Results

### *P.aurescens* TC1 genome scale metabolic network reconstruction

The genome of *P.aurescens* TC1 is composed of a circular chromosome of 4.6 Mb coding for 4,222 open reading frames (ORFs) as well as two plasmids, 0.3 Mb each, coding for additional ~600 ORFs^20^. To capture the most comprehensive metabolic capacities encoded in genome and plasmids, a genome-scale metabolic model was reconstructed. To this end, the genome and plasmids were annotated and used to create a draft reconstruction using the Model SEED pipeline^46^. Additional genome annotations of *P.aurescens* TC1 from four established and comprehensive public resources and databases (RAST, KEGG, JGI and UniProt) were then added to the draft network. Model Reactions involved in atrazine degradation pathway and its further conversion into the cellular building blocks isopropylamine and ethylamine were added manually (Fig. 1) based on a detailed reports of the pathway in the specific strain^18,47^. *P.aurescens* TC1 degrades atrazine by hydrolytic dechlorination. The process is catalyzed by triazine hydrolase (TrzN) followed by two hydrolytic deamination reactions catalyzed by hydroxyatrazine hydrolase (AtzB) and N-isopropylammelide isopropylaminohydrolase (AtzC) (reactions rxn06933, rxn03845 and rxn03846 in Fig. 1, respectively). These enzymes convert atrazine to cyanuric acid and release the alkylamines ethylamine and isopropylamine. Cyanuric acid accumulates stoichiometrically, whereas the alkylamines are further catabolized into cellular building blocks^21^. Ethylamine and isopropylamine are converted into acetaldehyde and L-alanine (Fig. 1), respectively, and can serve as sole sources of carbon and nitrogen required for growth ^18^. Relevant reactions that were missing in the automatic reconstruction were manually added to the model and are shown in Fig. 1. Simulations were carried iteratively to ensure the production of all biomass components under the minimal feasible conditions described in Strong, et al. ^18^. The reconstructed metabolic network presented here covers 25% of the ORFs present in the genome and includes 1176 genes, 2541 reactions, out of which 125 are exchange, and 2848 metabolites. The coverage of the genome is similar to those of previously published reconstructed metabolic networks of gram positive bacteria such as *C. difficile* or *B. subtilis* with 20% ^48,49^ and the overall number of reactions is similar to the number of reactions covered by gold standard, updated models of species with similar genome sizes such as *i*JO1366 ^50^. A detailed description of the network including the reactions, metabolites, genes, and compartments that comprise the network is provided in Supplemental data 1. The model is also available here as Systems Biology Markup Language (SBML) file ^51^ in Supplemental data 2. The SBML file can be used with tools such as MATLAB or other SBML compliant software. The minimal media used for simulations are available in Supplemental data 3.

**Figure 1.**
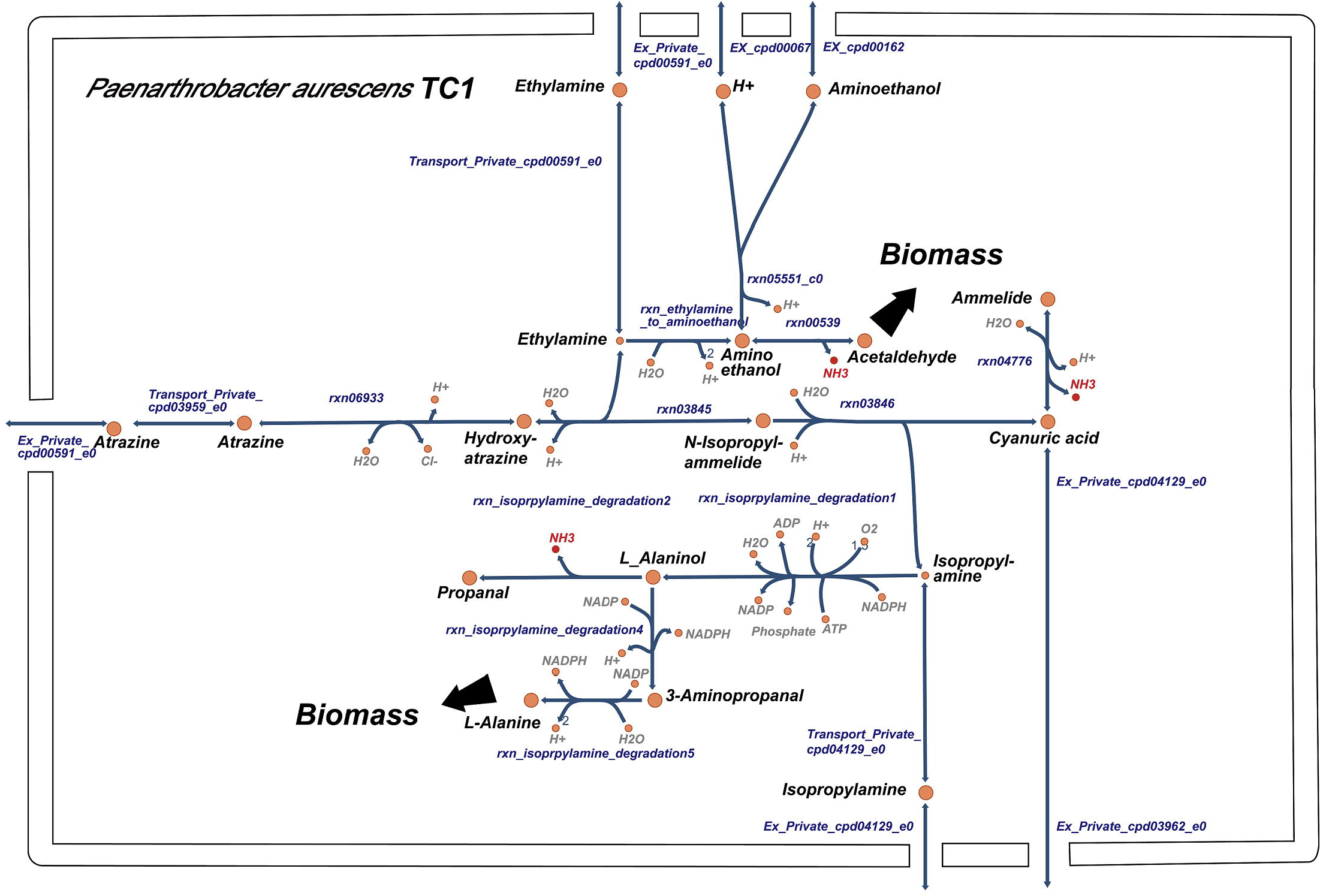
Atrazine metabolism as implemented in the cytoplasm compartment of *P. aurescens* TC1 genome scale metabolic model reconstruction (*i*RZ1176). External metabolites are notated with an e0;

### Validations of *i*RZ1176’s *in silico* predictions

As previously reported, *P.aurescens* TC1 degrades atrazine and metabolizes its intermediates, isopropylamine and ethylamine. Both can be used as sole carbon and nitrogen sources. Growth simulations confirm that the model captures these reported physiological capacities. Experimental growth measurements verify model simulations and show that atrazine and its degradation products isopropylamine and ethylamine can be used as sole carbon and nitrogen sources for *P. aurscens* TC1 (Fig. 2A). Supplementing the medium with glucose enhanced growth both *in silico* and *in vitro*, while the addition of ammonia had not shown an effect on growth. This is in accordance with previous work stating that *P.aurescens* TC1 growth is not inhibited by nitrogen sources such as ammonia ^22^, unlike the inhibition of atrazine degradation in *Pseudomonas sp. strain* ADP in nitrogen rich environments ^52^.

**Figure 2.**
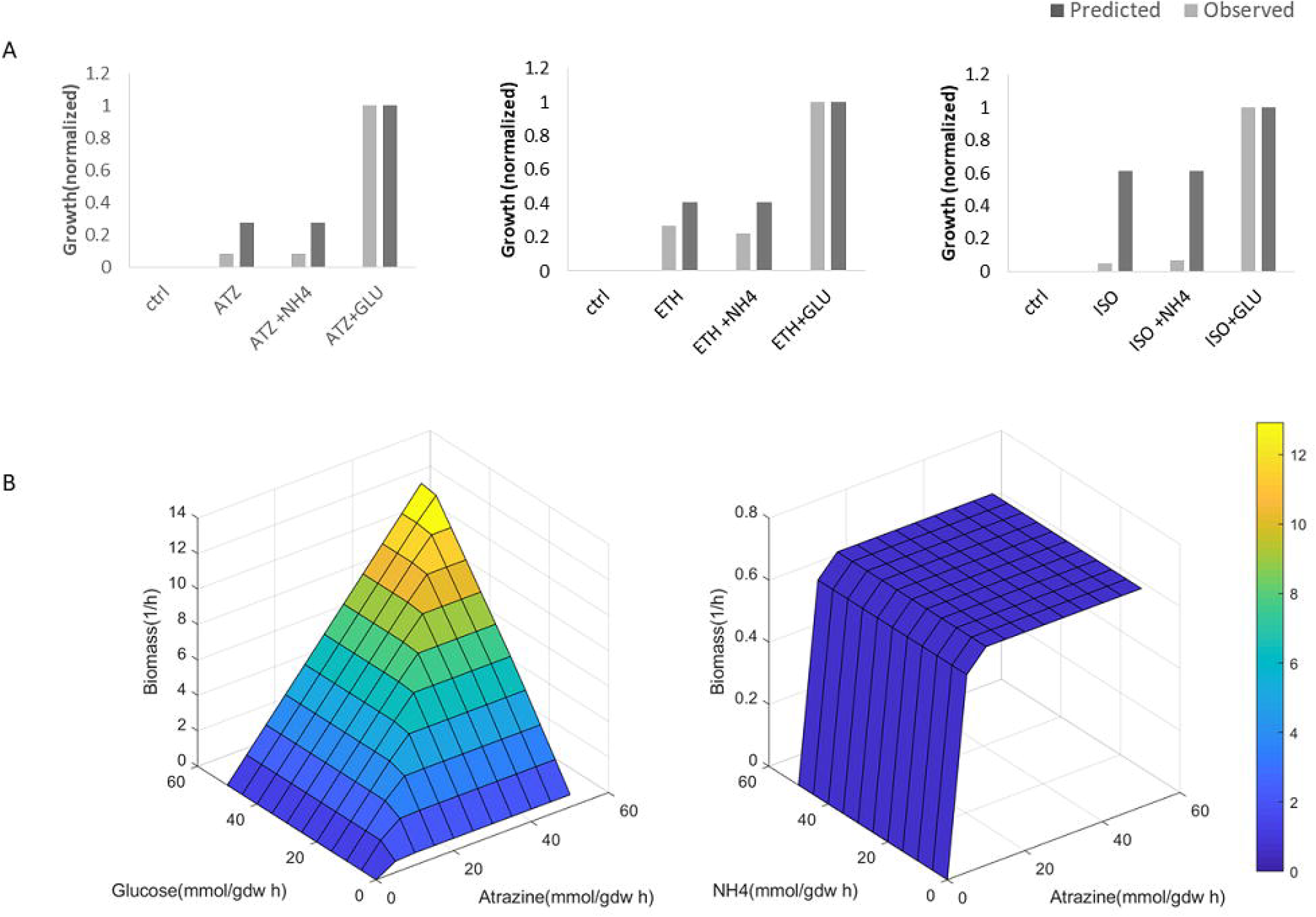
Growth performances of *P.aurescens* TC1 in dependence on the availability of different carbon and nitrogen sources. (A) Predicted and observed *A.aurescens* TC1 growth in mineral minimal media supplemented with Atrazine or its degradation intermediates. ATZ-Atrazine, NH4, GLU-Glucose, ETH - Ethylamine, ISO – Isopropylamine, CTRL - control. Values are normalized by dividing in maximal value. (B) Surface plot rendering *P.aurescens* TC1 growth (biomass) in dependence on atrazine, glucose and NH4 availability. Color scale is indicative of biomass (Z-axis).

Finally, growth simulations under a gradient of nitrogen *vs*. carbon sources further illustrate that atrazine serves as a good nitrogen source and as a poor carbon source (Fig. 2B). The growth promoting potential of combinations of atrazine with ammonium (NH_4_^+^) and glucose as nitrogen and/or carbon sources respectively was tested. The effect of nutrient ratio on growth rate was predicted by optimizing the production of biomass across uptake gradients while constraining the fluxes through exchange reactions for carbon and nitrogen sources. As expected, increasing glucose uptake increased the growth rate (Fig. 2B) and the maximal achievable growth rate was much higher with glucose as a carbon source in comparison to atrazine as a sole carbon source. However, atrazine is as equally good nitrogen source as ammonia. This may be endorsed by ammonium as a byproduct of the atrazine degradation process (Fig. 1).

### Media modifications effect on *P.* aurescens TC1 growth

In addition to glucose, several carbon sources were previously reported to enhance growth of several strains of *Arthrobacter* species in atrazine containing media ^21,22,53^. To identify additional media modifications that can potentially increase atrazine degradation, we simulated growth in 123 different media combinations, each supplementing the atrazine containing minimal mineral media with a single exchange metabolite. Several sugars, amino acids and dipeptides were predicted to induce enhanced growth (Supplemental data 4). This is in accordance with the previous characterization of *P.aurescens* TC1 as a metabolically versatile species ^20,21^. Out of a variety of potential nutritional sources, *Paenarthrobacter* species have been shown to utilize different amino acids in different capacities ^54,55^. The three amino acids isoleucine, histidine and methionine are predicted to support growth in high, mediate, and low efficiency, respectively (Supplemental data 4). Dynamic simulations of growth supplementing media with these three amino acids indicated that their effect is media dependent (Fig. 3). Isoleucine has the strongest predicted effect on growth in atrazine containing medium whereas histidine has stronger effect in the same medium without atrazine; methionine has the weakest effect in all media (Fig. 3). *P.aurescens* TC1 growth in *in vitro* media corresponding to the media used for the *in silico* simulations were generally in agreement with model’s predictions, supporting the superiority of isoleucine over histidine in atrazine supplemented media but not in the atrazine deficient medium (Fig. 3). Supplementing media with methionine showed a weaker effect on growth in comparison to the other two amino acids tested, agreeing with its weak effect in simulations. This is also in agreement with a recent work by Deutch ^55^ showing that whereas isoleucine and histidine serve as potential nitrogen and/or carbon sources of *P.aurescens* TC1, methionine does not serve as a growth supporter. Like atrazine, amino acids can potentially serve as both carbon and nitrogen source. Considering C:N ratio as an indicator of growth support potential, isoleucine and histidine have a ratio of 6:1 and 6:3, respectively, whereas methionine, having no growth promoting effect has a ratio of 5:1 (Supplemental data 4). Other amino acids with 5:1 or 5:3 C:N ratio like proline and serine, respectively, are also predicted to be an efficient supporters of growth, similar to isoleucine (Supplemental data 4). Thus, the growth effect of atrazine cannot be predicted merely based on C:N ratio.

**Figure 3.**
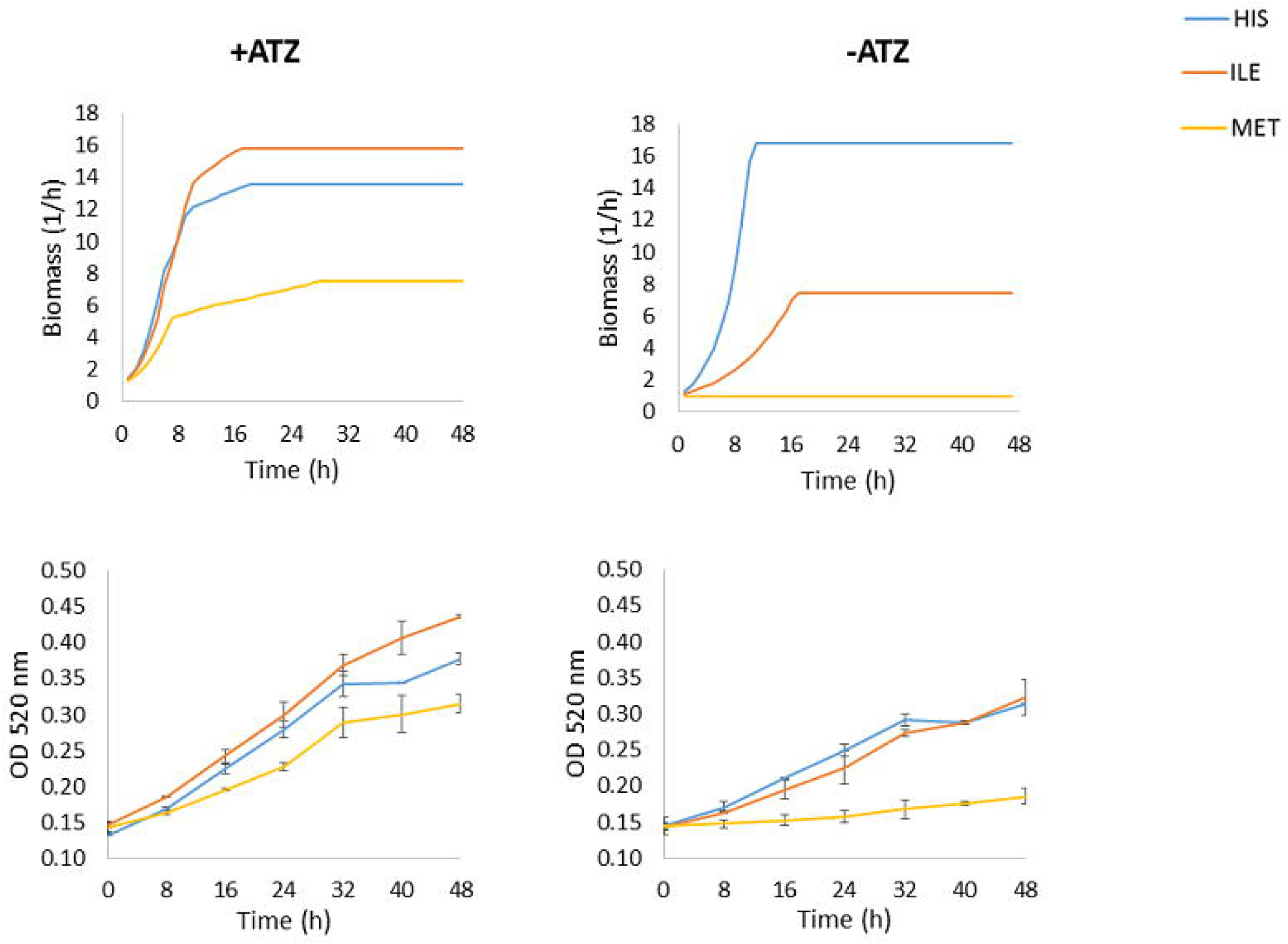
Predicted and observed growth performances of *P. aurescens* TC1 growth in mineral minimal media with glucose complemented by different amino acids and atrazine. ATZ-Atrazine, HIS-Histdine, ILE-Isoluecine and MET-Metionine. Observed values represents mean in triplicates; bars represent SD.

### Media modifications effect on atrazine degradation

Following the validation of the model predictions for growth capacities, we directly evaluated atrazine degradation in different media conditions. To this end, we applied Flux variability analysis (FVA), allowing the identification of the ranges of flux variability that are possible within a given media^56^. Once the biomass flux was maximized, we estimated the rate of atrazine degradation according to the relevant flux of interest (for example, phosphate). The simulation of atrazine degradation over time was in a medium that contains atrazine as a sole carbon and nitrogen source *vs*. modified media sets (see Methods). Atrazine degradation was estimated according to the amount of atrazine left in the media following each time cycle. Simulations showed that supplementing the media with glucose not only enhances growth (Fig. 2) but also induced an enhancement of atrazine consumption. Predictions were confirmed by experimental validation (Fig. 4).

**Figure 4.**
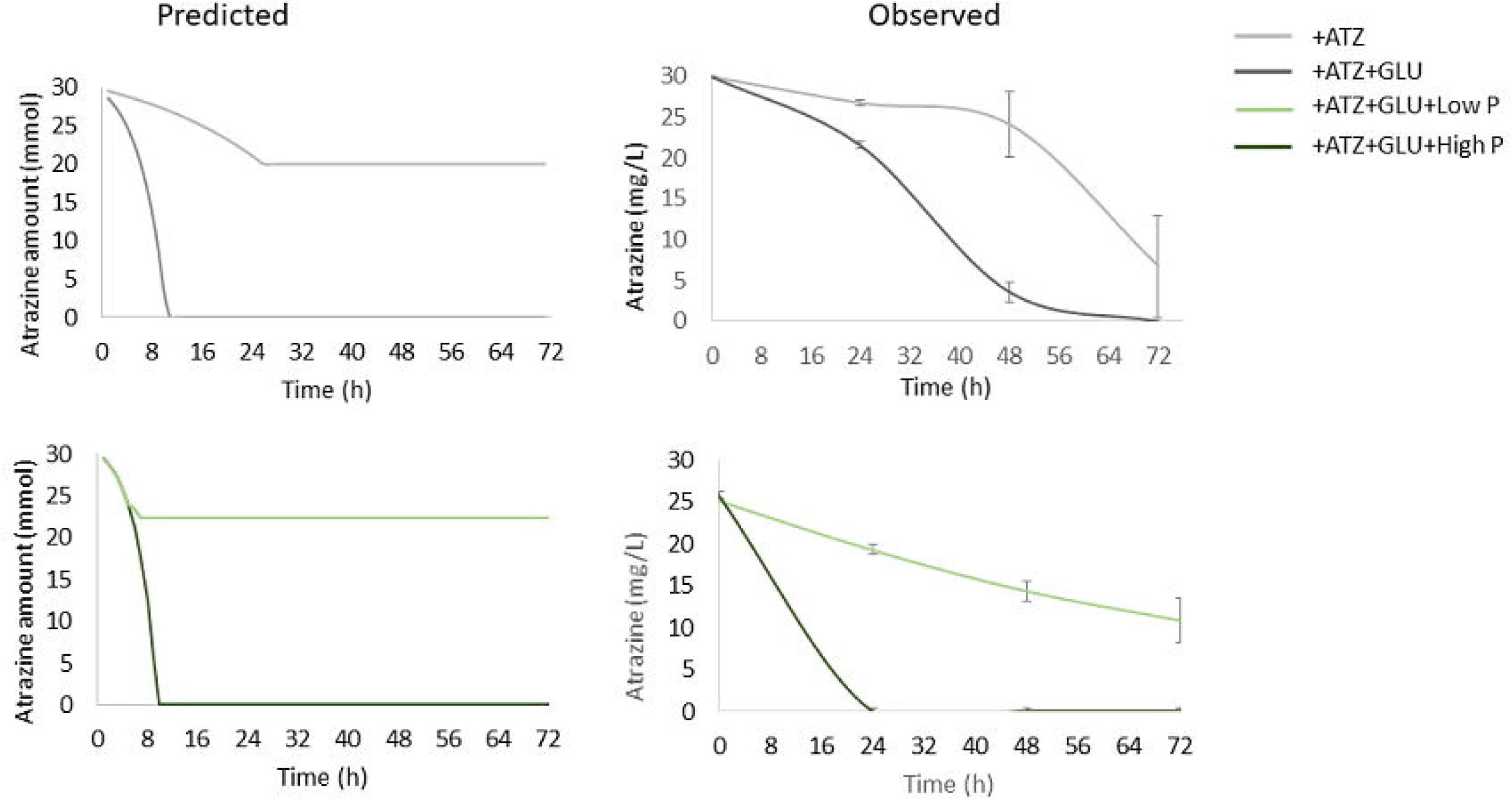
Predicted and observed atrazine degradation in modified media. High and low phosphate concentrations: 50 and 15 mmol/gdw h for predicted and 0.081 and 0.081 g/l for observed, respectively. ATZ-atrazine, GLU-glucose. Observed values represents mean in triplicates; bars represent SD.

Finally, the addition of phosphate, a main element in fertilizers, to soils has been reported to have a significant effect on the degradation of herbicides. For instance, its addition was shown to effect soil bacteria’s degradation activity including a reported effect on *Arthrobacter* sp. HB-5, a close relative of *P.aurescens* TC1^57^. Phosphate has been previously shown to accelerate atrazine degradation^3^. To test the effect of phosphate, we used the model to simulate growth and atrazine degradation in high or limited phosphate conditions. In accordance with these reports, simulations showed similar trends, which were confirmed by experimental results. These indicate an effect of phosphate concentrations *on P. aurescens* TC1 growth (Supplemental data 5) and atrazine degradation (Fig. 4).

## Discussion

The accumulation and persistence of herbicides and their intermediates in soil and drinking water has long been suggested to be disruptive, imposing a need for effective strategies for their degradation and removal ^58^. Bioremediation is a widely applied approach for the degradation of herbicides including atrazine ^9–13^. Here, we reconstructed a genome scale metabolic model, *i*RZ1176, of *P. aurescnes* TC1, a well-studied atrazine degrader, in order to use the *in silico* simulations for predicting optimal media to support its growth and atrazine degradation activity. The *in silico* simulations were carried on an array of minimal media using atrazine and its interim degradation products as alternative sole carbon and nitrogen sources supplemented by potential enhancers. The simulations suggested a group of metabolites promoting the increase in biomass yield including sugars, amino acids and dipeptides. We hypothesize that the introduction of such an efficient carbon source leads to an increase in biomass and a subsequent increase in the degradation activity. Simulations were confirmed by *in vitro* experimental validation. This genome scale reconstruction may be used for further experimenting and the gain of new biological knowledge of this metabolically versatile bacteria.

While the metabolic modeling approach presented here was demonstrated useful for generating predictions, its limitations should be acknowledged. First, model construction relies on automatic protocols and publically free genomic information. Biomass composition and energy maintenance requirements are hence approximations ^46,48^. Second, data used to construct the model is eclectic and annotations were gathered from numerous databases. The lack of data standardization has led to many overlaps and misnomers resulting in draft models containing massive gaps. The lion share of the gap filling and network improvement processes are still done manually due to the differences in data description and cataloging in the different databases. As technology and standardization advance, the process of constructing a metabolic model able to produce testable predictions will be faster and have greater accuracy. Despite relying on genomic data and a growth approximation, resulting predictions can successfully capture the relative effect of different media complements on growth and atrazine degradation. Accuracy of predictions is likely to be significantly improved by the addition of other layers of information including metabolomics, transcriptomics and reaction kinetics. These additional layers of information will allow imposing additional constrains on the internal and external fluxes, as well as an accurate description of the biomass composition specific to this strain. Continuation of the manual curation process, also based on added transcription information, is likely to improve the performances of future versions.

Overall, we describe the construction of a mathematical stoichiometric metabolic model of *Paenarthrobacter aurescens* TC1 (*i*RZ1176), allowing simulating its degradation activity. Simulations were used to predict growth and degradation following media modifications such as high/low phosphate content and the addition of variety of amino acids and carbon source to the standard minimal R medium. Effects of such media were observed experimentally to effect atrazine degradation. The simulations go beyond the C:N ratio of media components and take into account the complement of reactions and stoichiometric complexity of cellular processes and may be used to further study this versatile organism for the design of improved degradation strategies in soil. Additionally, we demonstrate that through using metabolic time dependent simulations, an optimal range of nutrient augmentation may also be planned. Though soil communities are complex in terms of their composition and microbial consortia dynamics, in a recent study we demonstrated the usefulness of modeling approaches for deciphering metabolic complexities^44^, pointing at the future potential of modeling approaches. Despite the complexity of soil communities, supplementing soil with carbon sources is long known to induce a change in microbial structure and function. A predictive tool such as the presented model is a first step towards designing optimal degradation efficiencies that can be further tested in the field. The integration of species-specific models together with community approaches is likely to contribute to our understanding of biodegradation processes in natural environments and promote the development of bioaugmentation and biostimulation strategies.

## Methods

### Reconstruction and curation of the *P. aurescens* TC1 metabolic model

An initial draft model of *P. aurescens* TC1 was reconstructed as described in Henry, et al. ^46^. Briefly, the complete genome sequence of *P. aurescens* TC1 was retrieved from NCBI (GenBank accession ids. NC_008711 including NC_008711.1 NC_008712.1, and NC_008713.1 – chromosome and two plasmids, respectively), uploaded into RAST ^59^, spliced into genes and annotated. RAST annotations were put through the Model SEED pipeline ^46^ and an initial metabolic network reconstruction was obtained. This initial reconstruction contained a list of reactions classified as exchange, transport, and cytosolic as well as a list of all relevant metabolites and a biomass reaction. Based on taxonomy, the gram positive biomass reaction was specified in the process of draft reconstructions using Model SEED and KBase pipelines. The composition of the biomass reaction summarizes the fractional contribution of a generalized microbial biomass precursors (e.g., amino acids and lipids) to the synthesis of a new cell and is similar to the previously published genome scale reconstruction of *B.Subtilis*^48^. Unspecified non-metabolic consuming processes in the cell were defined as ATP maintenance and set similarly to ^48^. Missing reactions were filled through an automatic gap-filling procedure, using default (optimal) media options ^46^. Manual curation procedures were carried to adapt model performances to reported physiological capacities. Simulation conditions were modified to reflect experimentally validated minimal media alternatives that support growth of *P.* aurescnes TC1 ^18^.

Curation procedures were carried iteratively versus growth simulations to ensure the production of all biomass components in Minimal Mineral Media (MMM: K^+^, Mn^2+^, CO2, Zn^2+^, SO_4_^−^, Cu^2+^, Ca^2+^, HPO_4_^2-^, Mg^2+^, Fe^2+^, Cl^−^), supplemented by alternative carbon and nitrogen sources including atrazine, ethylamine, isopropylamine, ammonia, glucose and selected amino acids. As part of the curation process, the hydrolytic pathway of atrazine degradation was *a priori* added to the model based on previously published biochemical evidence ^18^. Reaction directionality was assigned based on the KEGG database. Errors in reaction reversibility and elemental balance were manually identified and corrected according to literature and/or available databases. The model is available in SBML format and as excel sheet (Supplemental data 1 & 2). The minimal media used for simulations are available in Supplemental data 3.

### Flux balance analyses simulations and dynamic, time dependent simulations

Flux distributions in the metabolic model were determined using Flux Balance Analysis (FBA) ^60–62^. Briefly, a metabolic model includes a network that is described as a stoichiometric matrix **S**_*m* × *n*_, where *m* represents the number of metabolites and *n* the number of reactions in the model. In the assumed pseudo-steady-state, internal metabolites concentrations and fluxes are assumed to be constant. The model can be represented as: **S** × v = 0, where vector v signifies the fluxes through the internal reactions. As the number of reactions (*n*) typically exceeds the number of metabolites (*m*), the system is under-determined. In order to get a feasible solution space, linear programming (LP), subject to mass balance preservation and flux constraints, was used by introducing an optimization problem. Here, model growth was maximized by maximizing its biomass reaction, which was used as objective function. Specific fluxes for metabolites consumption and secretion were determined following fixation of the respective biomass reaction to its maximal value and then using Flux Variability Analysis (FVA)^56^, a linear programming method to find the minimal amount of metabolites needed to produce a preset amount of biomass. Dynamic modeling was used for the prediction of the profile of consuming metabolites typical to the biomass increase of *P. aurescens* TC1 and atrazine degradation across time. To this end, we simulated the behavior of our metabolic model across time. This was done as described in ^44^ and illustrated in Supplemental data 5, based on concepts defined in ^63^. Briefly, the model works under the following assumptions: (1) A finite start amount of media components is available; (2) A maximal amount of uptake a single cell can acquire from the media in a given time point is defined (the lower bound of the exchange reaction value); (3) New substrate concentrations in time t, are determined by the predicted substrate concentration for the previous step augmented with any additional substrates provided or consumed in the current iteration. The maximum uptake was set to a ratio of up to 1 unit of each metabolite available in the media; (4) After each time tick, the biomass amount was updated according to the flux amount of the biomass reaction in the model at this time tick. As biomass production rate serves as a proxy for the size of the population in the simulated environment and substrate uptake/secretion is mainly affected by to population size, the model was used to evaluate the actual substrate uptake/secretion and growth rate given the supplied media across time. Simulations were carried until reaching a state where time cycles did not lead to an increase in biomass. Initial concentration values for all metabolites were set to a fixed amount of 50 units (represented as the initial lower bounds, LB, of the exchange reactions). Simulation parameters and defaults are available in Supplemental data 3. Starting with one bacterium, the flux balance model was used to predict the uptake of phosphate, sulfate, carbon and nitrogen sources (*i.e.* atrazine and glucose) by *P. aurescens* TC1 across time.

All model simulations were done on an Intel i7 quad-core server with 32GB of memory, running Linux. The development programming language of our simulators was JAVA, and our linear programming software was IBM CPLEX. Surface plots were created using the MATLAB 2019a (MathWorks, Inc., Natick, Massachusetts, United States). Pathway maps were created using Escher maps^64^.

### *In vitro* bacterial growth conditions and chemical analysis

*P.aurescens* TC1 (ATCC BAA-1386™) was grown on R medium (based on ATCC 2662 R) plates with glucose (0.2%) and atrazine (30 mg/l, Agan Adama -Ashdod, Israel Israel) as sole nitrogen. Stock cultures were maintained in glycerol at −80°C. Cultures were resuscitated on R medium agar plates (1.5% Difco bacto™) and were used as starter colonies for liquid medium growth experiments. The growth of *P.aurescens* TC1 was tested in R media supplemented with different combinations of carbon and nitrogen sources, including Glucose (0.2%), Atrazine (30 mg/l, Agan Adama -Ashdod, Israel), ammonium chloride (37 mg/l), Ethylamine and Isopropylamine (31.1 and 40.8 mg/l, Sigma-Aldrich -Rehovot, Israel) and the amino acids Histidine, Isoleucine and Methionine (36, 92 and 103 mg/l respectively). Concentrations of amino acids, ethylamine and Isopropylamine were calculated to provide the same nitrogen content as the atrazine. In addition to the carbon and nitrogen sources, growth in media supplemented by atrazine and atrazine together with glucose was tested under two phosphate concentrations (K_2_HPO_4_ in 0.45 and 0.045 g/l). Each substrate was added to sterile R medium and bacteria was grown in triplicate 250ml flasks incubated in 30°C with shaking in the dark. One flask contained only R medium and was used as control. A 200 ul sample was taken from each flask every 8 hours for measurement of OD in microplate reader (520nm, Infinite^®^ 200 PRO Tecan, Männedorf Switzerland) and 1 ml was taken to study atrazine degradation rate using HPLC. The HPLC analysis was performed with Agilent 1100 HPLC (Waldbronn Germany) equipped with a DAD detector. Samples were separated on Kinetex C18 column (Phenomenex Torrance, CA) and the mobile phase consisted of 70% methanol flowing at 1 ml/min. Detection and quantitation of atrazine was done at 240 nm with detection limit of 0.1 mg/l. Total duration of each growth experiment was between 152 to 168 hours.

## Acknowledgments

This work was funded by grants from the NSFC-ISF joint program (31461143009), Israel Science Foundation Grant no. 1416/14. Atrazine was a gift of Agan Adama (Ashdod, Israel). We thank Omer Kapiluto for statistical support.

## Author Contributions

S.O, R.Z, T.L and S.F designed the study, constructed the model and performed *in silico* analysis. Z.R, S.P, R.A, and D.G designed and performed the *in vitro* experiments. Y.K and H.E analyzed and reviewed the results. All authors and approved the final version of the manuscript.

## Additional Information

### Supplementary information

Supplemental data 1 – Model reactions and metabolites

Supplemental data 2 – SBML model file

Supplemental data 3 – Simulation parameters and defaults.

Supplemental data 4 – Alternative media compositions

Supplemental data 5 – Supplemental figures

