## supplemental data files for "Genome-Scale reconstruction of *Paenarthrobacter aurescens* TC1 metabolic model towards the study of atrazine bioremediation": Supplemental_data5.pdf

**Supplemental data 5 - Genome-Scale reconstruction of *Paenarthrobacter aurescens* TC1 metabolic model towards the study of atrazine bioremediation**

Shany Ofaim[1,2]* †, Raphy Zarecki[1]*, Seema Porob[3], Daniella Gat[3], Tamar Lahav[1], Yechezkel Kashi[2], Radi Aly[1], Hanan Eizenberg[1], Zeev Ronen[3 †] & Shiri Freilich[1†]

[1]Newe Ya'ar Research Center, Agricultural Research Organization, Ramat Yishay, Israel, [2]Faculty of Biotechnology and Food Engineering, Technion-Israel Institute of Technology, Haifa, Israel, [3]Department of Environmental Hydrology & Microbiology, Zuckerberg Institute for Water Research, Jacob Blaustein Institutes for Desert Research, Ben-Gurion University of the Negev, Midreshet Ben-Gurion, 8499000, Israel

*equal contribution

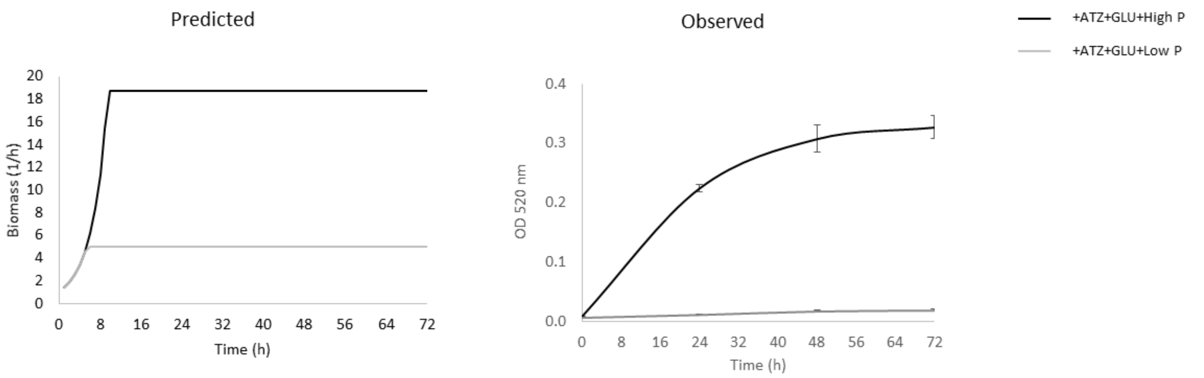


Figure S1 – *P.aurescens* TC1 growth in high and limited (low) phosphate conditions. High and low phosphate concentrations: 50 and 15 mmol/gdw h for predicted and 0.081 and 0.081 g/l for observed, respectively. ATZ- atrazine, GLU- glucose. Observed values represents mean in triplicates; bars represent SD.

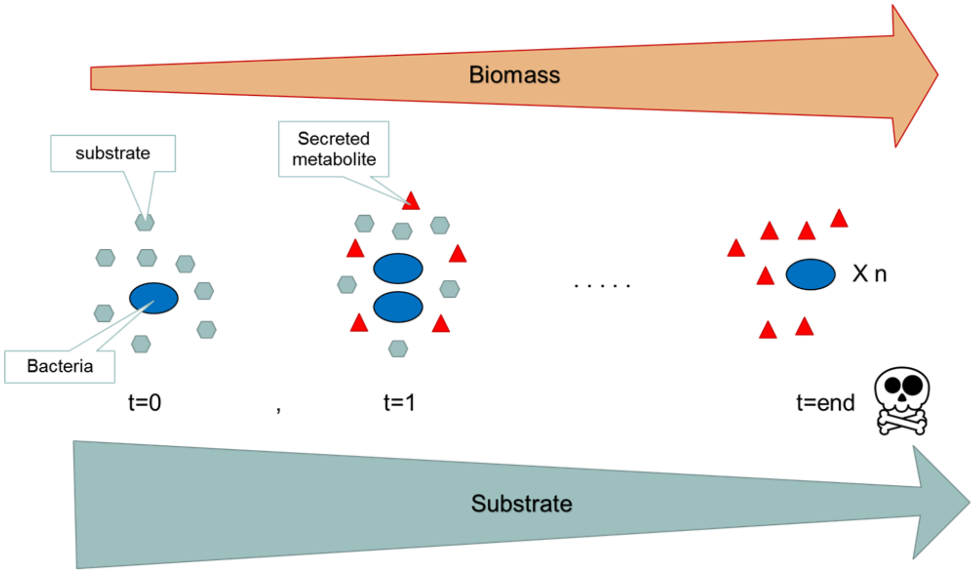


Figure S2 - A schematic diagram describing a time dependent simulation of nutrient dependent bacterial growth.
